# An outbreak of canine coronavirus type 2 in captive snow leopards *(Panthera uncia)* demonstrates a possible role for felids as mixing vessels for alphacoronaviruses

**DOI:** 10.1101/2024.03.25.586607

**Authors:** Ximena A. Olarte-Castillo, Abigail B. Schlecht, Paul P. Calle, Gary R. Whittaker

## Abstract

Coronaviruses are endemic and can cause disease in a wide range of domestic animal and wildlife species. The virus species *Alphacoronavirus-1* comprises a set of diverse viruses that are highly recombinogenic, including feline coronavirus type 2 (FCoV-2), which is a recombinant genotype of feline coronavirus type 1 (FCoV-1) and canine coronavirus type 2 (CCoV-2). Co-infection within a host promotes viral recombination; thus, to understand the origin of novel variants, it is crucial to identify hosts that can be infected with multiple alphacoronaviruses. The receptor for FCoV-2 and CCoV-2 is aminopeptidase N (APN), with the APN of the domestic cat *(Felis catus)* allowing entry of FCoV-2, CCoV-2, and other alphacoronaviruses. As wild felids are genetically closely related to the domestic cat, they may also be susceptible to these alphacoronaviruses. However, to date, natural infection with CCoV-2 has been reported exclusively in canids, not in felids. In this study, we retrospectively investigated a localized outbreak of enteritis in three captive snow leopards (*Panthera uncia)* at the Bronx Zoo (New York City, U.S.). Whole genome sequencing revealed shedding of CCoV-2 in the feces of the sick leopards. Phylogenetic analyses revealed it is related to highly pathogenic variants of CCoV-2 circulating in the U.S. and Europe. Comparative genetic analyses of the APN gene from five Asian wild felids, including the snow leopard, revealed a high percentage of identity to the APN of the domestic cat (>95.7%). These results emphasize the central role of domestic and wild felids in the emergence of recombinant alphacoronavirus.

## Introduction

Canine coronavirus type 2 (CCoV-2, *Alphacoronavirus*, *Coronaviridae* family) is a common virus of the domestic dog (*Canis lupus familiaris* (*1*)). Although normally considered to cause mild enteric disease, many pathogenic variants that cause severe gastrointestinal signs, have been reported in domestic dogs worldwide (2–5). Pathogenic variants of CCoV-2 that cause fatal systemic disease have also been described and are known as ‘pantropic’ CCoVs (pCCoVs (6)). The first pCCoV-2 variant reported (known as CB/05) was detected in Europe in 2005 (6), and pCCoV-2 CB/05-like variants have since been reported in several other European countries (7–9). Knowledge of CCoV- 2 in the U.S. is limited, but pathogenic non-systemic variants have been reported as the cause of fatal neonatal enteritis in domestic dogs (3). CCoV-2 can also infect wild species of canids (i.e. other members of the Canidae family (10)), including widespread urban species like the red fox *(Vulpes vulpes* (11)) and the raccoon dog (*Nyctereutes procyonoides* (*12*)). CCoV-2 can also infect wild species outside the Canidae family, including mustelids like the Chinese ferret-badger (*Melogale moschata* (*13*)), ursids (bears) like the giant panda *(Ailuropoda melanoleuca* (*14*)), and hyenids (hyenas), like the spotted hyena *(Crocuta crocuta* (*15*)). As with domestic dogs, CCoV-2 infection can result in subclinical signs or severe disease, including acute diarrhea (11, 16) and even neurological signs (14). A deadly outbreak of neurological CCoV-2 in captive giant pandas (14) highlights the potentially significant impact of CCoV-2 on endangered species. The recent detection of CCoV-2 in bamboo rats (*Rhizomys sinensis*) in farms in China (17) and humans with respiratory symptoms also emphasizes the need to investigate the host range and mechanisms for cross-species transmission of CCoV-2 (18, 19).

CCoV-2 belongs to the *Alphacoronavirus-1* species (20) together with canine coronavirus type 1 (CCoV-1), feline coronavirus type 1 and 2 (FCoV-1 and FCoV-2, respectively), transmissible gastroenteritis virus (TGEV), and porcine respiratory coronavirus (PRCV). Numerous recombinant variants among members of the *Alphacoronaviru*s-1 species have been documented (21). Variants of CCoV-2 that have recombined with TGEV circulate in Europe and are referred to as CCoV-2b.

(22, 23). Divergent variants resulting from the recombination between CCoV-1 and -2, known as CCoV-2c, have been detected in outbreaks in domestic dogs as early as 1976 in the U.S. (24), and recently in the UK (25). Additionally, the two CCoV-2s detected independently in humans also have signatures of recombination between CCoV-2 and FCoV-2 (18, 19, 26). Lastly, FCoV-2 is a recombinant genotype of FCoV-1 that acquired a region of its genome from CCoV-2 (27). Recently, a novel FCoV-2-like recombinant variant, FCoV-23, has spread among Cyprus’s feral/stray cat population, resulting in a 40-fold increase in reported FCoV-related deaths on the island (28). This outbreak shows that novel recombinant variants of FCoV-1 and CCoV-2 can emerge and rapidly spread in populations of free-ranging cats, posing a serious threat to animal and human health.

Host species susceptible to infection with several alphacoronaviruses, may promote recombination during co-infection events. CCoV-2, FCoV-2, TGEV, and PRCV use the aminopeptidase (APN) protein (also known as CD13) for host cell entry (29, 30). *In vitro* studies have shown that the APN of the domestic cat *(Felis catus)* (29–31) and the domestic dog (32, 33) can be used for cell entry by FCoV-2, CCoV-2, TGEV, and PRCV. Site-directed mutagenesis experiments have revealed the critical regions on the APN of the domestic cat (34) and the domestic dog (33) that are essential for binding to the receptor-binding domain (RBD) within the spike (S) protein of FCoV-2, CCoV-2, TGEV, and PRCV, The tertiary structure of the APN of the pig and the domestic dog coupled with PRCV (35) and CCoV-2, respectively (32), have shown that five residues are binding hot spots. Mutations in these sites can have adverse effects on virus-receptor interactions (34). Comparative genetic analyses of known protein receptors can help assess the susceptibility of various species to certain well-studied viruses, like SARS-CoV-2 (36). A recent study revealed that the APN of 14 African species of wild carnivores (including canids and felids) is highly conserved and similar to that of the domestic dog and cat (>92.7% pairwise similarity (15)). This suggests that wild felids, like domestic cats, may be susceptible to infection with FCoVs and other alphacoronaviruses, potentially playing a key role in the emergence of recombinant variants. However, the sequence of the APN gene of wild felids outside of Africa has not been obtained, and the genetic diversity of this receptor remains understudied. In order to determine whether wild felids can act as mixing vessels for alphacoronaviruses, it is important to conduct comparative genetic analyses of the APN protein as a first step to assess their susceptibility to these viruses. Additionally, although infection or exposure to FCoVs has been reported in some species of wild felids, to our knowledge, natural infection with CCoVs has not been documented in felids (including the domestic cat). Thus, it is also necessary to investigate whether FCoV and CCoV naturally infect and circulate in domestic and wild felids.

The snow leopard (*Panthera uncia*) is a wild felid found throughout the mountains of Central and Western Asia (37). Currently, this species is classified as vulnerable in the International Union for Conservation of Nature (IUCN) Red List (38). Due to their secretive nature, there is limited information on infectious diseases that may affect this species (39). However, reports from captive animals have shown that, like other wild felids (40, 41), the snow leopard is susceptible to infectious diseases common in domestic cats, including feline panleukopenia virus (42) and FCoV (43). Lethal viruses common in wild and domestic canids, like canine distemper virus (42, 44), and human viruses, like SARS-CoV-2 (45), have also been reported in captive snow leopards. Although free-ranging leopards may not be greatly impacted by infectious diseases due to their solitary lifestyle in isolated areas (39), the expansion of livestock herding into these remote areas (46) may increase their risk of exposure to infectious diseases from domestic animals and livestock. Thus, understanding the impact of common infectious diseases of domestic cats and dogs on snow leopards could aid conservation efforts for wild populations.

Here, we report a retrospective molecular epidemiological investigation of a localized outbreak of diarrhea that occurred in 2012 in captive snow leopards at the Bronx Zoo in New York City, U.S. Pancoronavirus screening detected CCoV-2 in the fecal samples of the three sick leopards. To confirm the identity of the virus, we sequenced the whole genome using next-generation sequencing. Genetic analysis of the whole genome did not find any recombination breakpoints, confirming it is a CCoV-2 and not an FCoV-2. Phylogenetic analyses indicated the detected CCoV-2 is closely related to pathogenic variants reported in the U.S from 2009 and 2013, and pCCoV-2 reported in Europe. To assess the genetic susceptibility to CCoV-2 and FCoV-2 of Asian wild felids, we obtained the complete sequence of the APN gene of the snow leopard, the Pallas’ cat *(Otocolobus manul),* the leopard cat (*Prionailurus bengalensis*), and the Malayan and Amur tiger (*Panthera tigris altaica* and *jacksoni*, respectively). Comparative genetic analyses revealed a high genetic identity to the sequence of the APN of the domestic cat (>95.7%), with all the substitutions observed in essential residues being conservative. This study provides the first genetic evidence of CCoV-2 infection in a wild felid, resulting in gastrointestinal signs. This study also predicts that FCoV-2, CCoV-2, TGEV, and PRCV may be able to infect various wild felid species due to the high genetic conservation of the APN gene within the Felidae. These results highlight the importance of felids (wild and domestic) as mixing vessels for the emergence of recombinant alphacoronaviruses and the necessity of conducting surveillance of both FCoV and CCoV in canids and felids.

## Methods

### Sample collection

At the end of January 2012, three snow leopards at the Bronx Zoo started exhibiting signs of gastroenteritis. Two of the individuals, a four-year-old male (PU293) and a two-year-old female (PU294), experienced intractable diarrhea and vomiting, and a third one, an eleven-year-old female (PU296), experienced diarrhea and anorexia. All three leopards were born and raised at the Bronx Zoo. Feces were collected from the three individuals, and an additional vomit sample was taken from PU294. After collection, samples were frozen at -80°C until further use. After treatment with fluids and parenteral antibiotics, all three snow leopards recovered.

To sequence the APN gene, additional kidney samples were collected from four Asian felid species that died at the Bronx Zoo from unrelated causes. Kidney samples were collected from a leopard cat in 2009, a Malayan tiger in 2019, a snow leopard in 2021, and an Amur tiger in 2024. An additional intestine sample was collected from a Pallas’ cat that died of feline infectious peritonitis in 2008 in a different zoological institution in the U.S. (40). After collection, samples were frozen at -80°C until further use.

### RNA extraction from fecal, vomit, and tissue samples

Approximately 40mg of feces and vomit from the three sick leopards or tissue (kidney or intestine) from five Asian wild felids were used for RNA extraction using the Monarch^®^ Total RNA Miniprep Kit (New England Biolabs, NEB) according to manufacturer’s instructions and including the suggested DNase I in-column step. Before the extraction, the samples were homogenized on a Bead Blaster 24 (Benchmark) bioruptor using 600 μl of lysis buffer and 500mg of ∼2.3mm Zirconia/Silica beads (Biospec Products) for the vomit and tissues, and ∼0.1 mm zirconia/silica beads (Biospec Products) for the fecal samples. cDNA was synthesized using the LunaScript®RT SuperMix Kit (NEB) following manufacturer’s instructions. The cDNA of the feces and vomit samples was used for coronavirus (CoV) screening, and the cDNA from the tissues of the five Asian wild felids was used to amplify the APN gene.

### Screening, whole genome sequencing, and genetic analyses of CCoV-2

Screening for CoV was carried out using previously described pancoronavirus primers targeting a conserved region of the RNA-dependent RNA-polymerase (RdRp) gene (47). The resulting PCR products were sequenced in a MinION Mk1B (Oxford Nanopore Technologies, ONT) using the Native Barcoding Kit 24 V 14 (ONT) and a Flongle Flow Cell (R10.4.1, ONT). The sequence of the whole genome of the detected CCoV-2 was obtained using the previously described primers (6) to amplify the 3’-end of the genome (including the structural and accessory genes), and a new set of primers (Table S1), to amplify the 5’-end of the genome. The primers designed in this study (Table S1) were pooled into two pools (1 and 2) to do a total of two multiplex PCRs. The size of all the amplicons is ∼2,000bp. The Q5 High-Fidelity 2X Master Mix (NEB) was used to carry the multiplex PCRs.The resulting PCR products were sequenced as mentioned above for the screening but using a Flow Cell R10.4.1 (ONT). The resulting sequences (from the screening and the whole genome sequencing) were duplex basecalled using Dorado v0.5.1 (ONT). Highly accurate reads were mapped against the sequence of CCoV-2 HLJ-071 (accession number KY063616) using BWA-MEM2 v 2.2.1+galaxy1 (48, 49). The three whole-genome sequences obtained in this study (28,511 nt) were aligned with 103 additional whole-genomes, including 35 TGEV, 29 FCoV-1, 1 CCoV-1, 3 FCoV-2, and 34 CCoV-2. This and the additional 4 alignments mentioned below were performed using the MUSCLE algorithm (50). This alignment was used to detect recombination breakpoints in the obtained sequences using the Recombination Detection Program (RDP (51)) method in RDP-5 (52). An additional alignment of the whole genome sequences obtained in this study and CCoV-2 S378/1978/USA (KC175341), pCCoV-2 CB/05/2005/Italy (K981644), FCoV-1 RM/2002/USA (FJ938051), and FCoV-2 NTU156/2007/Taiwan (GQ152141), was used to generate a similarity plot using SimPlot 5.1 (53).

Three additional alignments of different genomic regions were performed. One included the complete open reading frame 1a (ORF1a) region of the genome (11,902 nt) of 33 CCoV-2, 4 FCoV-1, 3 FCoV-2, 1 TGEV, and 1 CCoV-1. The second and third included a partial region of the S1 domain (the N-terminal domain, NTD, 212 nt), and the S2 domain of the S gene (224 nt), respectively. These two alignments included 70 CCoV-2 and 4 FCoV-2 sequences and were performed to include the only available sequences of CCoV-2 variants circulating in domestic dogs in the U.S. between 2009 and 2013 (3). Each of these alignments was used to compute the best-fitting nucleotide substitution model and to construct a maximum likelihood (ML) phylogenetic tree using MEGA 11 (54). Statistical support of the tree branches was estimated by performing a bootstrap with 1,000 replicates. The trees were visualized using iTOL v6 (55). The complete amino acid sequence of the S protein of the three CCoV-2s obtained in the study was aligned with 2 TGEV, 5 FCoV-2, and 58 CCoV-2 to calculate the pairwise percentage similarity in Geneious Prime 2023.2.1.

### Sequencing and genetic analyses of the APN gene of wild Asian felid species

The sequence of the complete APN gene (2,748 nucleotides) was obtained from five Asian wild species using the primers listed in Table S2. The primers were pooled into two pools (1 and 2), to do two multiplex PCRs. The size of all the amplicons is ∼900bp. The PCRs were carried out using the Q5 High-Fidelity 2X Master Mix (NEB) and sequenced and assembled as mentioned above for the whole genome sequencing of CCoV-2. The five sequences obtained in this study and 17 additional sequences available in GenBank of other wild and domestic species were translated to their respective amino acid sequences. The 17 other sequences included the APN of the domestic pig *(Sus scrofa),* 8 canids, including the African wild dog *(Lycaon pictus),* the dingo *(Canis lupus dingo),* the silver-backed jackal *(Canis mesomelas)*, the bat-eared fox *(Otocyon megalotis),* the artic fox *(Vulpes lagopus),* the red fox, the raccoon dog, and the domestic dog, and 8 felids, including the fishing cat *(Prionailurus viverrinus),* the jaguarundi *(Puma yagouaroundi)*, the bobcat *(Lynx rufus),* the clouded leopard *(Neofelis nebulosa),* the Geoffroy’s cat *(Leopardus geoffroyi),* the cheetah *(Acinonyx jubatus)*, the African lion (*Panthera leo)*, and the domestic cat. The 22 translated sequences were alignment using the Clustal Omega algorithm (56) in Geneious Prime 2023.2.1 (Biomatters Ltd). Based on this alignment, a Maximum likelihood tree was constructed in MEGA 11 (54). This alignment was also used to calculate the percentage of amino acid identity of the complete sequence of the APN and to assess the genetic variation of the 23 residues known to interact with the virus directly (32, 34, 35). When substitutions were observed in the sequence of the wild species when compared to the domestic cat or dog sequences, each was classified as conservative, semi-conservative, and non-conservative when there was a substitution with an amino acid with similar biochemical properties, weakly similar biochemical properties, and no similar biochemical properties, respectively.

## Data availability

The sequences of the three whole-genomes of CCoV-2 and the complete APN genes from five Asian wild felids obtained in this study were deposited in GenBank under accession numbers 7006 – 7013.

## Results

In total, four samples, including the feces from three leopards and the vomit of one of the leopards, were screened for CoV RNA. The feces from the three snow leopards (PU293, PU294, PU296) were positive, and the vomit of PU294 was negative. The whole genome sequences of the positive samples were obtained, and they were 28,511nt long and 99.9% similar. The three genomes contained 11 ORFs, including ORF1a, ORF1b, S, ORF3a, ORF3b, ORF3c, Envelope (E), Mat ix (M), Nucleocapsid (N), ORF7a, and ORF7b. An initial phylogenetic tree of the complete ORF1a (11,902 nt) placed the three coronaviruses detected in the CCoV-1/CCoV-2 group and not in the FCoV-1/FCoV-2 group (Figure 1).

**Figure 1.**
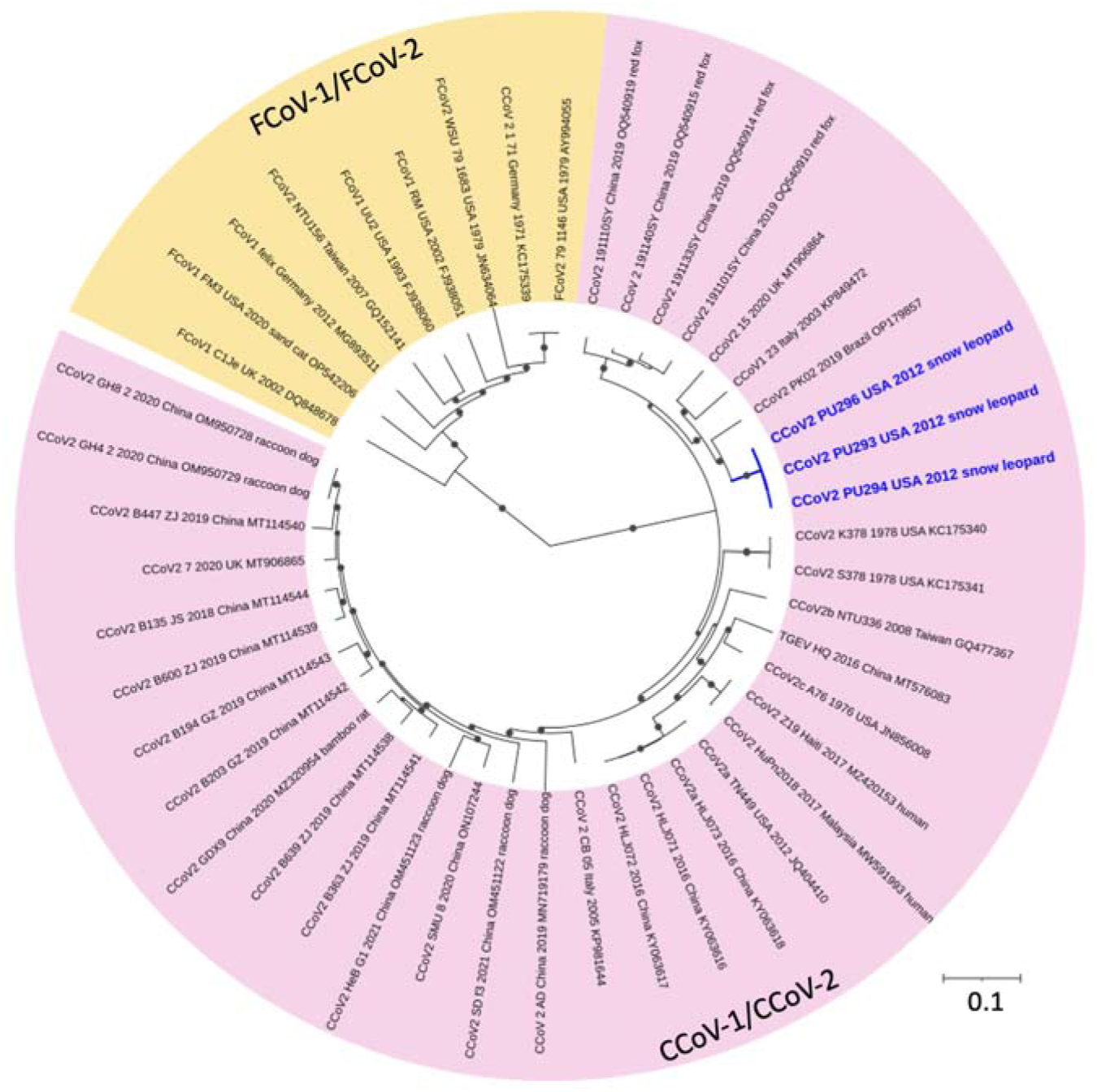
Maximum likelihood phylogenetic tree of the complete ORF1a (11,902nt) of selected FCoVs and CCoVs. Two groups can be observed, one including FCoV-1 and FCoV-2 (in yellow) and another one including CCoV-1 and CCoV-2 (in pink). In dark blue are the three sequences obtained in this study, which group together within the CCoV-1/CCoV-2 group (in pink). For all variants included, the sequence name includes the type of CoV (FCoV-1, -2, CCoV-1, -2), year and place of collection, and accession number. For variants recovered from wild animals, the species’ common name is after the accession number. Dots in the branches indicate a bootstrap value higher than 0.6. The estimated best-fitted nucleotide substitution model was HKY+I+G.

To explore if the CoVs obtained in this study were recombinant (i.e. FCoV-2, CCoV-2b, CCoV-2c) or not (i.e. CCoV-2), their whole genome sequences were compared to 103 sequences of other alphacoronaviruses, including TGEV, FCoV-1, CCoV-1, FCoV-2, and CCoV-2. No deletions, insertions, or recombination breakpoints were detected in the three genomes compared to the other alphacoronaviruses included in the analysis. A genetic comparison of the whole genome of PU294 and an FCoV-2 (NTU156/2007/Taiwan), an FCoV-1 (RM/2002/USA), and pCCoV-2 CB/05/2005/Italy showed that along the whole genome, PU294 is consistently more similar to CCoV-2 and not to FCoV-1 (Figure 2). In contrast, the recombinant portion of FCoV-2 NTU156/2007/Taiwan, which includes a partial region at the 3’-end of ORF1b to a partial region at the 5’-en of the N gene (Figure 2), is more similar to CCoV-2 than FCoV-1, indicating its CCoV-2 origin.

**Figure 2.**
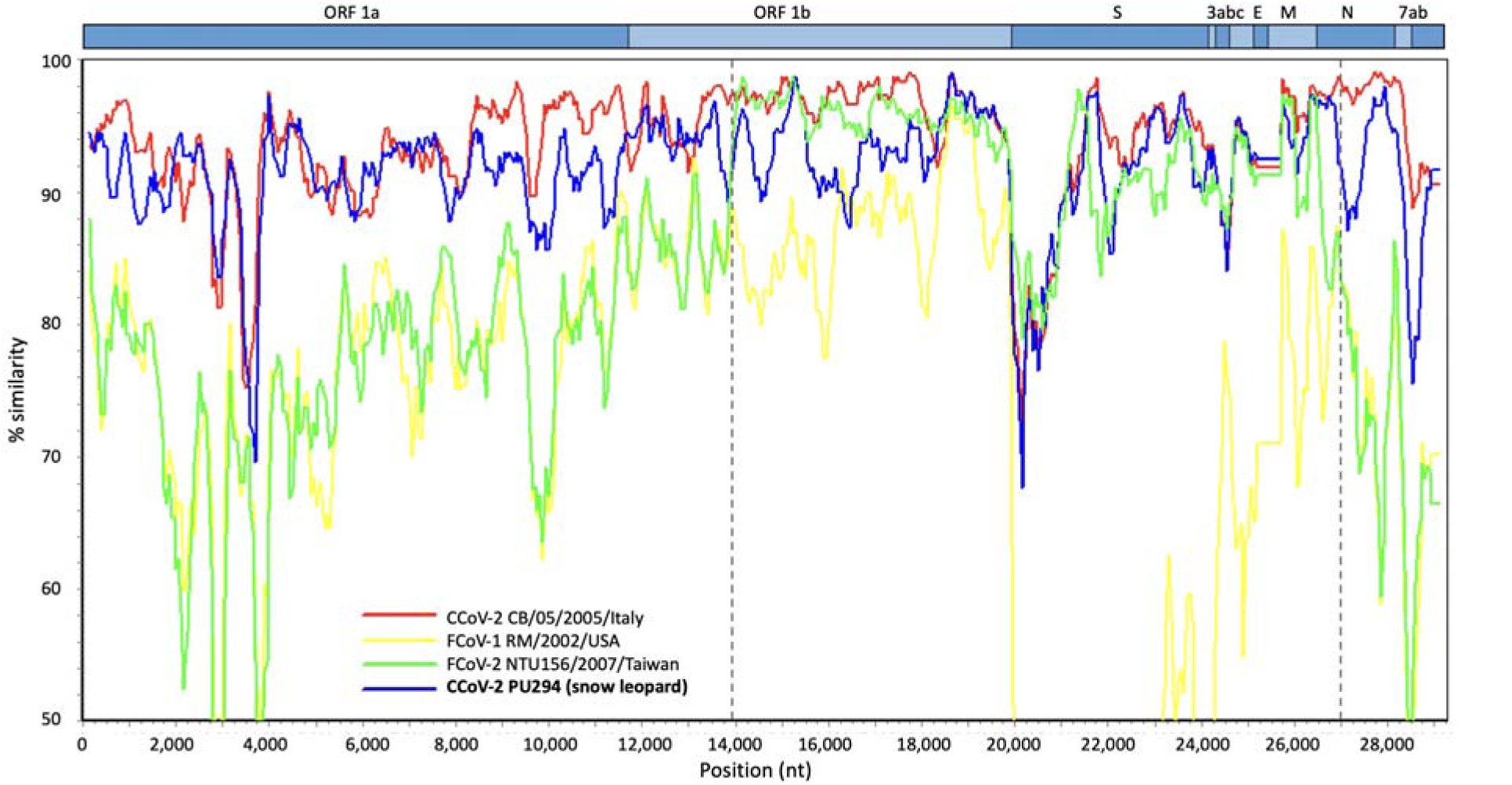
Similarity plot comparing the complete genome sequence of the CCoV-2 detected in one of the snow leopards (PU294) with other canine and feline coronaviruses. A graphical representation of the genome above the chart shows the location of each gene. The genome sequence of CCoV-2 S378/1978/USA (KC175341) was used as a reference. In red is CCoV-2 CB/05/2005/Italy (K981644), in yellow is FCoV-1 RM/2002/USA (FJ938051), in green is FCoV-2 NTU156/2007/Taiwan (GQ152141), and in dark blue is one of the CCoV-2 detected in a snow leopard (PU294). Dashed vertical lines indicate the recombination breakpoints of FCoV-2 NTU156/2007/Taiwan (in green). Within this region, FCoV-2 is more similar to the CCoV-2 variants (in dark blue and red) than to FCoV-1 (in yellow). In the rest of the genome, FCoV-2 is more similar to FCoV-1 (in yellow). The CCoV-2 from the snow leopard (in blue) is highly similar throughout the whole genome to the reference CCoV-2 and CCoV-2 CB/05/2005/Italy (in red), indicating that it is a CCoV-2 and not an FCoV-2. The graph was constructed using the Kimura 2 parameter distance model with a window of 300bp and a 50bp step.

Two additional phylogenetic trees of partial regions of the S gene (S1 and S2 domains, Figure 3A and 3B, respectively), were constructed to compare our three sequences to other CCoV-2 circulating in the U.S. during the period when the outbreak in the snow leopards occurred. In these phylogenies, the three sequences from the snow leopards obtained in 2012 grouped together within a group containing other CCoV-2 from domestic dogs in the U.S obtained from 2009 to 2013 (in dark pink in Figure 3), as well as the pCCoV-2 variants CB/05/2005/Italy and 450/2007/Italy (in green in Figure 3) in the phylogeny of the S2 domain (Figure 3B), and NA/09/2009/Greece in the phylogeny of the S1 domain (Figure 3A). These phylogenetic trees also included CCoV-2 variants from different wild species, including red foxes and raccoon dogs detected in China (indicated in Figure 3 with a square and a triangle, respectively). For the CCoV-2 reported in wild animals, at least 4 groups are circulating in China. The pairwise similarity of the amino acid sequence of the complete S protein of the three sequences obtained in this study was 99.8% similar. When compared to 65 additional sequences of CCoV-2, FCoV-2, and TGEV, the highest percentage similarity of the S protein of the snow leopard CCoV-2 was with pCCoV-2 CB/05/2005/Italy (95.6%), followed by pCCoV-2 450/2007/Italy (95.5%).

**Figure 3.**
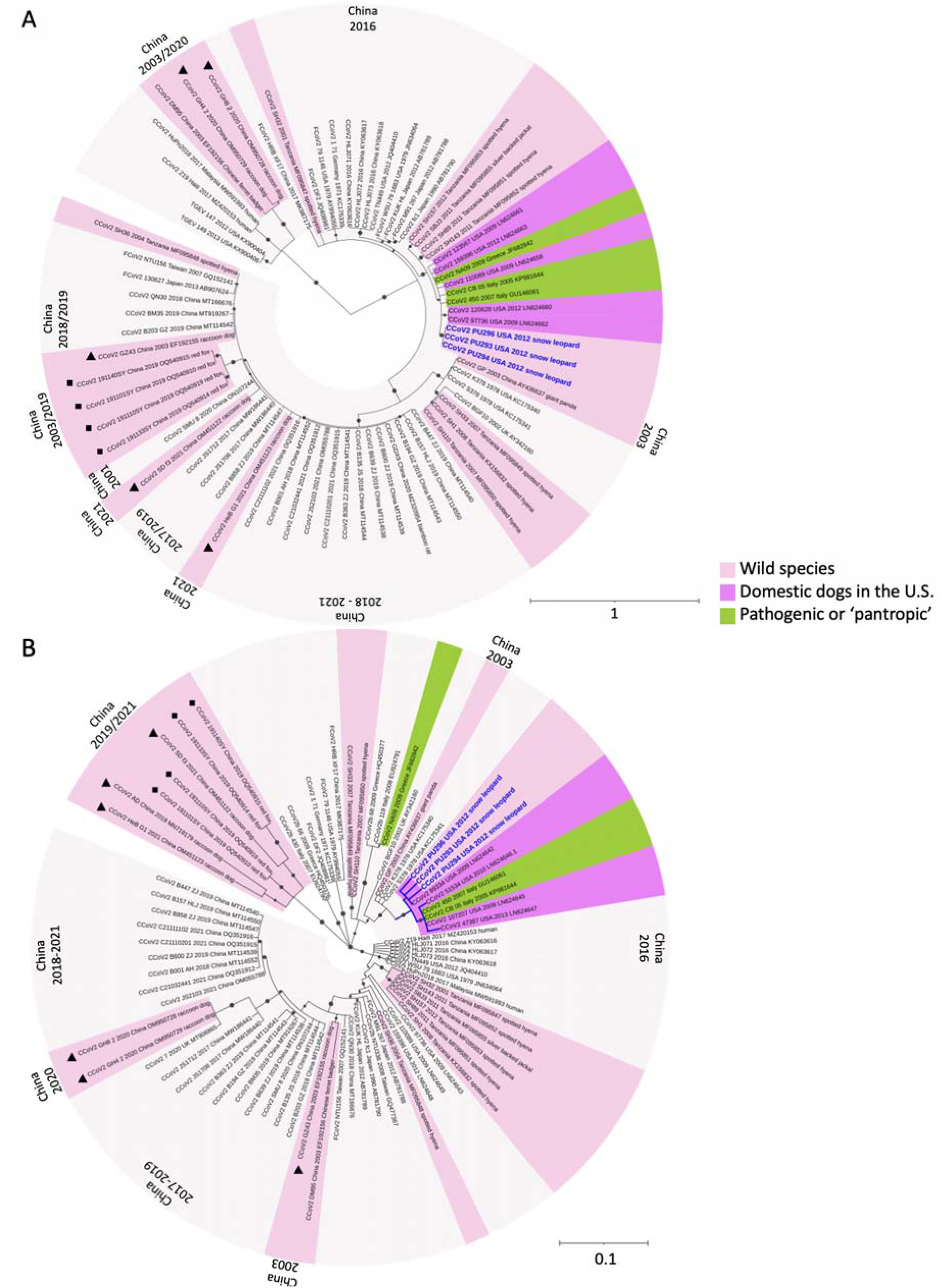
Maximum likelihood phylogenetic tree of a partial region of (A) the S1 domain (212nt) and (B) the S2 domain (224 nt) of S of selected FCoV-2 and CCoV-2. The three sequences obtained in this study are in dark blue. In light pink are the variants recovered from wild species, in light gray are those from domestic dogs, in dark pink are those from domestic dogs in the U.S., and in green are the pathogenic or ‘pantropic’ CCoV-2 variants from domestic dogs. The variants detected in China (from domestic or wild species) are labeled with the location and year of collection at the tip of the tree. From the variants detected in wild animals in China, those from red foxes are indicated with a square, and those from raccoon dogs are indicated with a triangle. In both trees, the three sequences obtained in this study (in blue) group together with other sequences of CCoV-2 from domestic dogs collected in the U.S. in 2009, 2010, 2012, and 2013 (in dark pink) and two to three pathogenic or ‘pantropic’ CCoV-2 variants (in green). For all variants in both trees, the sequence name includes the type of CoV (FCoV-2, CCoV-2), year and place of collection, and accession number. The species’ common name for variants recovered from wild animals is after the accession number. Dots in the branches indicate a bootstrap value higher than 0.6. The estimated best-fitted nucleotide substitution model for both alignments was GTR+I+G.

We obtained the complete APN gene sequence from five wild Asian felids, including the snow leopard, to compare their genetic similarity with the APN of the domestic cat, dog, and pig, in order to assess their susceptibility to the alphacoronaviruses of these domestic species. A phylogenetic tree of the complete APN protein sequence revealed the grouping of the sequences in the two expected groups (Canidae and Felidae, Figure 4). Within the Felidae, the domestic cat was grouped with the bobcat, and the snow leopard was in a different group together with three other Asian wild felids, including the Amur and Malayan tiger and the clouded leopard. A comparison between the amino acid sequence of the domestic cat and the 12 wild felids included in the tree revealed a high amino acid identity ranging from 95.7% with the African lion to 98.8% with the bobcat (Figure 4). The sequence of the snow leopard was 96.8% identical to that of the domestic cat (Figure 4). Within the Canidae, the domestic dog grouped together with the Silver-backed jackal. The two species of canids that have home ranges in Asia, the raccoon dog and the red fox, were placed in a different group together with the arctic fox and the bat-eared fox (Figure 4). A comparison between the amino acid sequence of the domestic dog and the 7 other wild canids included in the tree revealed a high amino acid identity ranging from 97.0% with the red fox to 99.1% with the silver-backed jackal (Figure 4). A comparison between the Canidae and Felidae revealed 90.5% amino acid identity and 89.3% when comparing felids and canids with the domestic pig.

**Figure 4.**
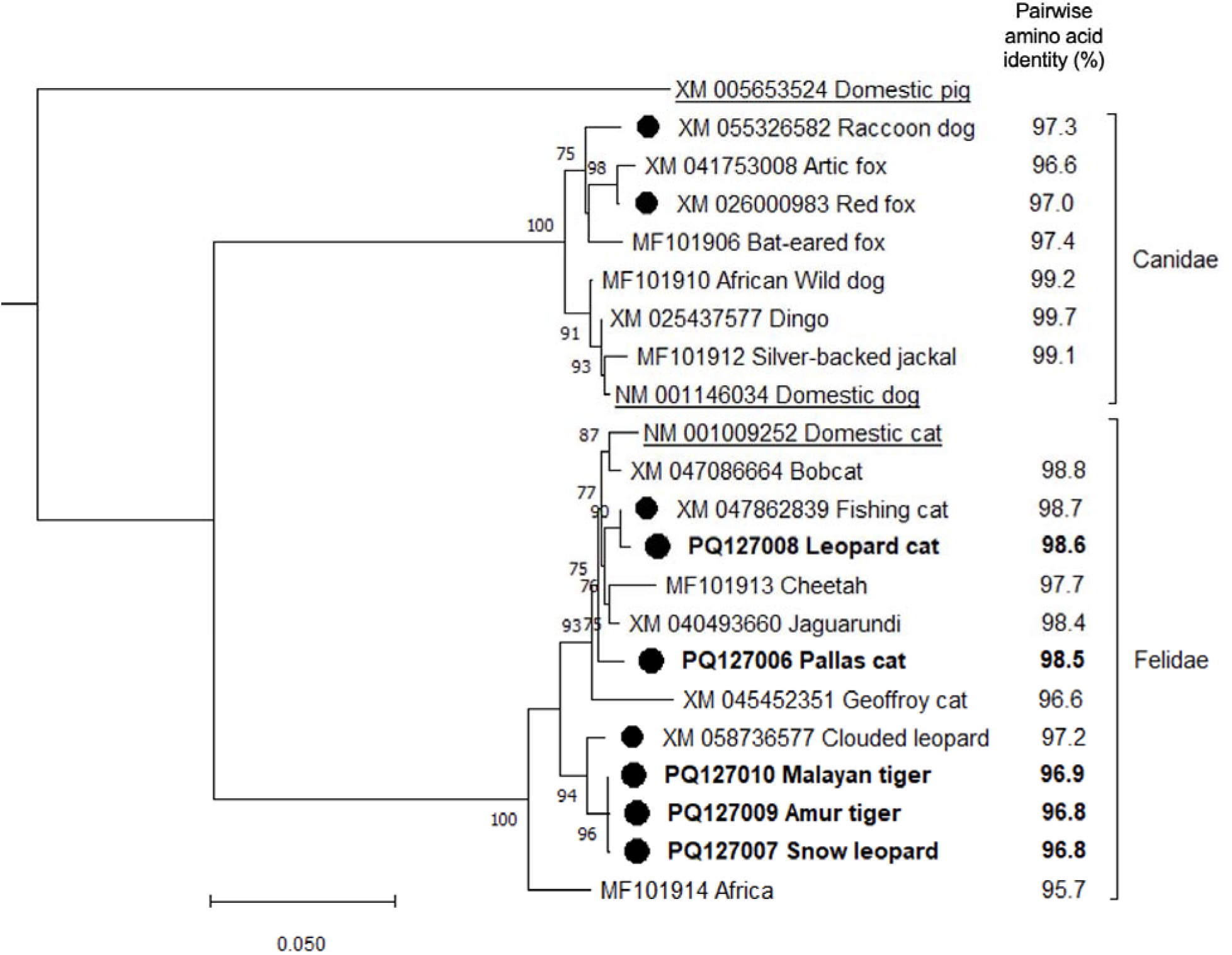
Maximum likelihood phylogenetic tree of the complete amino acid sequence of the APN from 13 felids, 8 canids, and the domestic pig. The five sequences obtained in this study are in bold. The sequences from the domestic cat, dog, and pig are underlined. Species of wild canids and felids with a home range in Asia are marked with a black circle. Next to the common name of each species is the value of the percentage of the pairwise amino acid identity between the domestic dog and the other wild canids within the Canidae and the domestic cat and the other wild felids within the Felidae. Numbers in the branches indicate bootstrap values for 1,000 replicates. Only bootstrap values hig er than 75 are shown.

A comparison of the 23 residues in APN known to interact with CCoV-2 and PRCV between the domestic cat and the 12 wild felids included in this study revealed that these residues are highly conserved (Table 1). There were substitutions in a maximum of three residues (371, 768, 779). However, all were conservative (i.e., the substituted amino acid has similar biochemical properties). Moreover, the substitutions of two of these three residues (768 and 779) in the snow leopard, the Malayan and Amur tiger, and the African lion, were identical to those observed in the canids included in the analysis (E768K and Q779E). Overall, when comparing these 23 residues between the 21 felids and canids included in the alignment, a maximum of seven substitutions were observed. Of these,5 were conservative (371, 739, 768, 779, 790), one was semi-conservative (771), and one was not conservative (370). When comparing the residues of felids and canids with those of the pig, there was only one additional substitution (743), which was conservative. (Table 1).

**Table 1.**
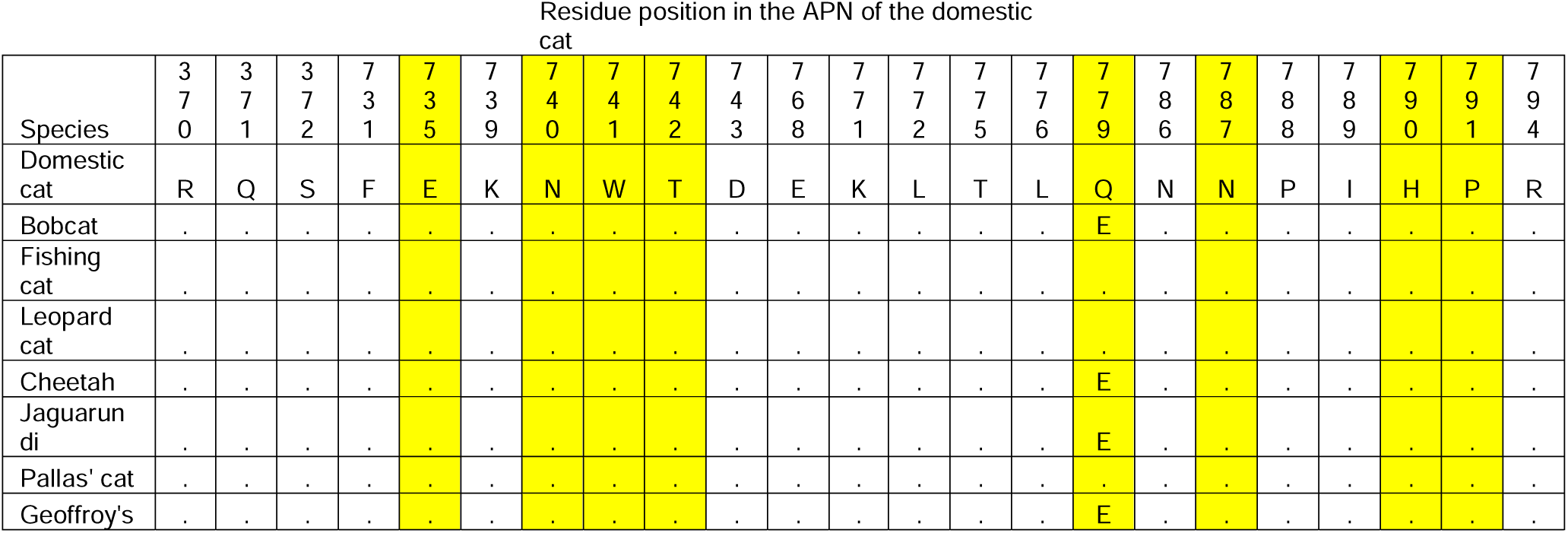
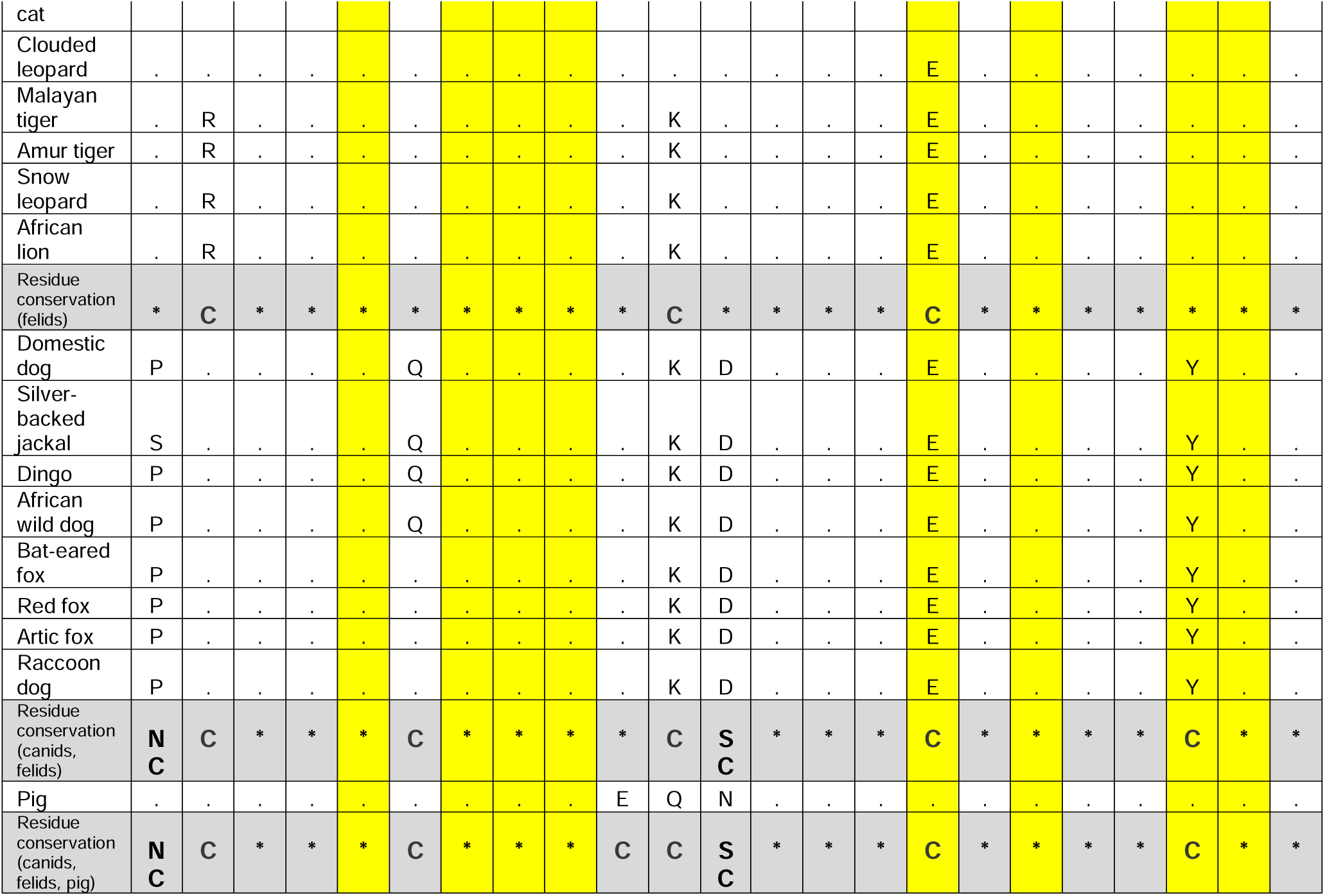
Genetic variation of the 23 residues in APN that are essential for the interaction with the virus in 22 species of felids, canids, and the domestic pig. The sequence of the domestic cat was used as reference. A dot indicates a matching residue with the reference sequence, and residues that differ are explicitly shown. Shaded in gray is the residue conservation scores of each comparison. An asterisk (*) indicates all residues were identical, C represents conservative substitutions, SC represents semi-conservative substitutions, and NC represents non-conservative substitutions. Shaded in yellow are the five residues essential for viral entry. Four of these (735, 740, 741, 742) are conserved in all the species included in the table.

## Discussion

Recombination is a significant mechanism that shapes the genetic diversity of coronaviruses, allowing the emergence of new variants with altered pathogenicity and immunogenicity (57, 58). For example, FCoV-2 is a recombinant genotype of FCoV-1 in which the S gene (and flanking regions) has been exchanged with that of CCoV-2. As the S gene of FCoV-1 and CCoV-2 is highly divergent (<54% pairwise amino acid identity (21)), their antigenic (59) and host cell entry properties (60, 61) are vastly different. The diverse recombinant variants of FCoV-2 reported to date display different recombination breakpoints (27, 62, 63), indicating multiple recombination events between FCoV-1 and CCoV-2. The emergent FCoV-23 is also a novel recombinant variant in which FCoV-1 most likely acquired its S gene from a highly pathogenic CCoV-2 (28). Recombinant alphacoronaviruses also circulate in domestic dogs (22). These recombinant variants have also exchanged different regions of the S gene with other alphacoronaviruses, like TGEV (CCoV-2b recombinants (23)) and CCoV-1 (CCoV-2c recombinants (24)). Therefore, identifying which species are susceptible to infection with both FCoVs and CCoVs is crucial for understanding which hosts may facilitate co-infections and thus promote recombination. Early experimental infection studies have indicated that domestic cats can be infected with CCoV-2, with variable levels of viral shedding in feces (64). Additional *in vitro* studies have shown that the APN of the domestic cat supports the infection of several alphacoronaviruses, including FCoV-2, CCoV-2, TGEV, PRCV, and even the human CoV-229E (HCoV-229-E (29–31)). Furthermore, persistent infection of FCoV-1 is common in domestic cats (1), which may increase the chances of co-infection with other alphacoronaviruses. The recent report of the cross-species transmission of FCoV-1 between a domestic cat and a captive wild felid (a Pallas’ cat), shows that wild felids may also be susceptible to FCoV-1 (40). This experimental and empirical evidence suggests that domestic cats likely play a key role in the origin of recombinant variants of alphacoronaviruses. In this study, we conducted a retrospective genetic analysis of a localized outbreak of diarrhea and vomiting in three captive snow leopards. Analysis of whole genome sequences (Figure 2) and phylogenetic data (Figure 1) revealed the presence of CCoV-2 in the feces of the three sick snow leopards. These results support the idea that, as experimentally shown in domestic cats (64), other felids can be infected with CCoV-2. This underscores the importance of domestic and wild felids as hosts that can be infected with various closely related alphacoronaviruses, which may promote recombination.

The binding interaction between virus and their host receptor is one of the most important determinants of CoV host range and cross-species infection (65). In this study, we show that the APN of the snow leopard and five other Asian wild felids is highly similar to that of the domestic cat (96.8% to 98.7% pairwise amino acid identity, Figure 4). When compared to the sequence of the domestic cat, we found substitutions in 3 of the 23 residues in APN essential for viral entry, in the snow leopard and the Amur and Malayan tiger sequences (Table 1). However, all three were conservative substitutions, and two of these (768, 779) have the same residue as those observed in canids (Table 1), including the red fox and the raccoon dog. In the other Asian wild felids included in the analysis, including the fishing cat, Pallas’ cat, and leopard cat, the 23 residues were identical to those of the domestic cat. Therefore, we predict that these species may be highly susceptible to FCoV-2 and CCoV-2 (and perhaps TGEV and PRCV) infection due to their high genetic similarity to the domestic cat in a protein crucial for the virus-host interaction. Our findings of CCoV-2 in three snow leopards (Figure 1), together with previous reports of FCoV infection in this species (43), are consistent with this hypothesis. These results yield important insights into the potential susceptibility of Asian wild felids, including endangered species like the Malayan (critically endangered) and Amur tiger (endangered) to alphacoronaviruses (15).

Although CCoV commonly causes subclinical to mild gastrointestinal signs, specific mutant or recombinant variants can be highly pathogenic. The S protein of CoVs binds to the host cell receptor and triggers the fusion with the host cell membrane (66). Thus, it is an essential determinant of cellular tropism and pathogenicity. The phylogenetic trees of the partial S1 and S2 domains (Figure 3); placed the three variants obtained in this study together with other variants of CCoV-2 detected in domestic dogs in the U.S. between 2009 and 2013 (in dark pink in Figure 3), from which only partial genomic sequences are available (3). These CCoV-2 variants were associated with severe enteritis (3), consistent with the gastrointestinal signs observed in the three snow leopards. However, the genetic comparison of the amino acid sequence of the S protein showed that the highest overall similarity (>95.5%) was with two pCCoV-2 variants (CB/05/2005/Italy and 450/2007/Italy). The phylogenetic trees of the partial S1 and S2 domains (Figure 3) also showed that the three CCoV-2 obtained in this study were closely related to these pCCoV-2 variants (CB/05/2005/Italy and 450/2007/Italy, in green in Figure 3). pCCoV-2 has been reported in several European countries and is linked to fatal outbreaks in pups in which viral systemic infection was detected (i.e., CCoV-2 RNA was detected in lungs, spleen, liver, kidneys, mesenteric lymph node, and brain). In contrast, although the CCoV-2 variants in the U.S. were also fatal, the virus was confined to the intestines (3). Further studies are necessary to comprehend the molecular mechanisms behind the extended tropism of pCCoV-2 compared to closely related strains with restricted tropism reported in the U.S., including the ones in this study. It is worth noting that all the pathogenic CCoV-2 variants reported in the U.S. were from neonate dogs (3), whereas the three leopards in which we found the related CCoV-2 were all adults older than 2 years. The S gene of FCoV-23 is highly similar (>97% pairwise nucleotide similarity) to other pCCoV-2, NA/09/2009/Greece (28). Thus, the deadly outbreak in cats in Cyprus could be the result of the emergence of a recombinant FCoV that obtained the S from a highly pathogenic CCoV-2. The results of this study suggest that variants of CCoV-2 highly similar to highly pathogenic variants circulating in domestic dogs can infect a felid species, the snow leopard, causing gastrointestinal signs. If these CCoV-2 can infect felids, recombination with FCoV-1 could occur, and the exchange of S can result in recombinant variants with distinct tropism, pathogenicity, and host range. Therefore, it is necessary to assess if both FCoVs and CCoVs naturally circulate in wild and domestic felids and canids. It is essential to use next-generation sequencing techniques to sequence entire genomes for detecting co-infections in a single individual and evaluating the probability of the emergence of recombinant variants in a given population.

Studying the potential impact of infectious diseases on free-ranging populations of snow leopards is challenging due to their secretive nature (39). However, infectious diseases may have a great negative impact on juveniles and neonates, as shown for captive snow leopards (67). A previous study found FCoV infection and shedding in healthy snow leopards in two zoological institutions in the U.S. (43). In contrast, we report that CCoV-2 infection can result in gastrointestinal signs and viral shedding in feces. Due to their solitary lifestyle, snow leopards may acquire pathogens from other carnivore species (wild or domestic (39)). China is estimated to be home to more than 50% of the world’s snow leopard population, making it a crucial figure for the conservation of this species (68). China’s economic growth is causing human encroachment on the habitat of snow leopards (46, 68) which may lead to their increased exposure to pathogens of domestic and wild urban species. This study shows that at least four groups of CCoV-2 are circulating in wild species in China and at least 3 in domestic dogs (Figure 3). Of epidemiological interest are the CCoV-2s circulating in common wild urban species. For example, a high prevalence of CCoV-2 was detected in urban raccoon dogs at the beginning of the SARS-CoV-2 pandemic (12). These abundant species may interact with other wild and domestic animals in urban areas, potentially increasing the risk of cross-species virus transmission and recombination. This is especially concerning given the high prevalence of CCoV-2 in domestic dogs in mainland China (69). An additional risk for snow leopards and other wild species are the highly pathogenic variants of CCoV-2 that have been reported in several deadly outbreaks in China (14), including the recent outbreak of a divergent CCoV-2, which has been linked to the death of over 39,000 red foxes at a rescue center in the Guizhou province in China (11). Likewise, co-infection with CCoV-1 and CCoV-2 was reported in red foxes and raccoon dogs sampled in China in 2005 (70), which may promote the emergence of novel recombinant variants in abundant wild species. Surveillance of both FCoV and CCoV in domestic and wild carnivores in China is essential to understanding the risk of cross-species transmission to vulnerable species, like the snow leopard.

## Acknowledgments

Work in the Whittaker lab is funded in part by the Cornell Feline Health Center. We thank Bonnie Raphael and Dr. D McAloose for collecting the samples of the sick snow leopards and the additional tissues, respectively.

