## Supplementary material for "An outbreak of canine coronavirus type 2 in captive snow leopards *(Panthera uncia)* demonstrates a possible role for felids as mixing vessels for alphacoronaviruses": Table S1, Table S2

**Table S1.** Primers used for the amplification of the CCoV-2 detected in the three sick snow leopards. The primers were pooled into pools 1 and 2, to do two multiplex PCRs. The size of all the amplicons is ~2,000bp.

| Primer ID | Primer sequence (5’–3’) | Primer pool |
| --- | --- | --- |
| 1_F_CCoV2 | TTCTTACCGAACCCTCCGTCAT | 1 |
| 1_R_CCoV2 | CGAGCMGCATTACAGAGCTTR | 1 |
| 2_F_CCoV2 | CATGCTGGTGATGTTGAAAATGTCT | 2 |
| 2_R_CCoV2 | CCTTCAAGTTCGAYACCYTGGT | 2 |
| 3_F_CCoV2 | ATGGGTGGTGGTGACAAAACTG | 1 |
| 3_R_CCoV2 | AAGGCAGCAGCATCATGRAAA | 1 |
| 4_F_CCoV2 | AATGTYAACCATGARCGTGTGTC | 2 |
| 4_R_CCoV2 | ACCTCTTGACTACTGTACTTCTTGTG | 2 |
| 5_F_CCoV2 | TGCAAGGCAGACACGTATYCC | 1 |
| 5_R_CCoV2 | ATGGAGCCATRTTAATRTAAGCAAT | 1 |
| 6_F_CCoV2 | TTGCWTGCCTGCTATTGCA | 2 |
| 6_R_CCoV2 | AGCATGTTYACAAAGCAGAAAGC | 2 |
| 7_F_CCoV2 | TGCCTGTKGACATGCAAGGT | 1 |
| 7_R_CCoV2 | GCRCCACCATAAGARTCTTGTTGTG | 1 |
| 8_F_CCoV2 | GTTCTTGGTTAYATCGGYGCAACAG | 2 |
| 8_R_CCoV2 | AGTGCATCCTGCTCTTCATATGA | 2 |
| 9_F_CCoV2 | GGTGARGCTGCTATGACAGACTT | 1 |
| 9_R_CCoV2 | ACYGAATTTCTGTTAAGTGGRGGCT | 1 |
| 10_F_CCoV2 | TCGGYTCAGRGGATGTTGAAGA | 2 |
| 10_R_CCoV2 | TGTAAGYTCAAGACCACCAGCC | 2 |
| 11_F_CCoV2 | GTTGTAAAAGCACGAGCACCAC | 1 |
| 11_R_CCoV2 | ACAACAACTTTTCARTGTACTGTCAG | 1 |
| 12_F_CCoV2 | CTCTYTAYAGAGCGTATGTTGARGA | 2 |
| 12_R_CCoV2 | AGTAATCCAACRACCATTTCATTCAA | 2 |
| 13_F_CCoV2 | CACTTGGTGGATCTGCTGC | 1 |
| 13_R_CCoV2 | CCAGTGGTAGTTGCTGTTGT | 1 |

**Table S2.** Primers used for the amplification of the complete APN gene from five Asian wild felids. The primers were pooled into pools 1 and 2, to do two multiplex PCRs. The size of all the amplicons is ~900bp.

| Primer ID | Primer sequence (5’–3’) | Primer pool |
| --- | --- | --- |
| APN_1F | ATCCTGGCCATCCTCCTGG | 1 |
| APN_1R | AGATCCGGATCAGGACGCC | 1 |
| APN2_F | CCGAGTTCGAAACCACACCC | 2 |
| APN2_R | CAAGGAGGAAGTGCTGCTGG | 2 |
| APN3_F | GTGAGTGCCATCATGGACCG | 1 |
| APN3_R | GTCAGCCTCGTTCACCAGTTC | 1 |
| APN4_F | ACAACCCGATCCACCCCAA | 2 |
| APN4_R | ATGGGAAGGTGAGAGAGGGC | 2 |
